# Identifying factors that may improve mechanistic forecasting models for influenza

**DOI:** 10.1101/172817

**Authors:** Pete Riley, Michal Ben-Nun, James Turtle, Jon A. Linker, David P. Bacon, Steven Riley

## Abstract

Influenza causes substantial morbidity and mortality and places strain on healthcare systems, some of which could be mitigated by accurate forecasting. Specific humidity and school vacations have both been shown independently to affect the transmission dynamics of influenza at large spatial scales. Here, we compare the ability of five compartmental transmission models, which include these two processes, to explain influenza-like-illness (ILI) incidence data for five United States counties for which school vacations and specific humidity data were available over a span of four seasons. We used the models in two different ways. First we fitted all available data at the same time and assessed model performance using standard measures of parsimony and goodness-of-fit. Then we conducted a retrospective forecasting study in which we attempted to predict incidence beyond a given week by fitting to data available up to that week. In general, when fitting the data using the whole season, we found that either specific humidity, school closures, or a combined model incorporating both effects captured the variability in incidence better than a fully constrained SIR-like model. Moreover, where these factors play a role, the timing of the variations suggests a causal relationship. When school vacations and specific humidity were important, the model-estimated parameters were broadly consistent. Retrospective forecasting simulations were consistent with the explanatory use of the models, with both specific humidity and school vacations giving more accurate forecasts than a simple SIR-like model in some populations and for some seasons. Our results suggest that influenza forecast models should test for the importance of different factors such as school vacations and specific humidity on a population-by-population and year-by-year basis.

**Author summary:** Understanding the underlying factors that contribute to the transmission of influenza is crucial for developing models with predictive capabilities. In this study, we address two key effects: humidity and school vacations. We show that both can play an important role, depending on the location of the population as well as the timing of the school vacations. We then demonstrate how such mechanistic models can be used to forecast an influenza season as it unfolds, including estimates of the uncertainty of the predictions at each week of the forecast. To make better influenza forecasts, models need to test whether factors such as school vacations and specific humidity are important for a given season and population.

## Introduction

Mechanistic models of infectious diseases [1] are frequently used during outbreaks of emerging human and animal infections to forecast key features of epidemic curves. For example, during the 2001 foot-and-mouth outbreak in the UK, models were used to forecast the rate of incidence decline and the total number of farms affected when culling response time was decreased [2, 3]. In 2009 in Singapore, real-time forecasts of ILI rates were given for the first time prospectively online [4] and also communicated privately in real-time to policy makers in a number of locations. Real-time prospective forecasts have also been made for the Ebola outbreak in west Africa in 2014/15 [5] and for the Zika outbreak in the Americas in 2015 [6].

Outside of outbreaks, in temperate climates, seasonal influenza epidemics are also a forecasting target [7–9]. Because there is considerable variation from year to year in the amplitude of the peak, its timing, and the total number of epidemic weeks, influenza can present potential resource allocation issues for clinical management teams and public health officials. In some years, during peak incidence of ILI, respiratory health services can be overwhelmed and intensive care units saturated, with “knock-on” effects to other parts of the the healthcare system [10, 11].

Models of influenza incidence, whether used for forecasting or as retrospective epidemiological tools, have been applied at different spatial scales and have included a large variety of mechanisms and methodologies. For example, the impact of school closures on transmission has been modeled for cities, such as Hong Kong [12], and Countries, such as France [13], while the contribution of climatic drivers has been assessed for U.S. states [14] as well as individual cities [8].

Meteorological conditions have long been thought to play a role in the transmission dynamics of influenza [15]. Heuristically at least, this is supported by the radically different evolutions of ILI profiles in tropical versus temperate zones [16]. Theoretically, if at least some of the virus transmission is airborne [17], support for such a relationship comes from the idea that the effective “lifetime” of the virus in a droplet is sensitive to the local conditions within which it is embedded. Experimentally, it has been shown that the transmission rate amongst guinea pig hosts increased as the relative humidity decreased [18].

In this study, we describe a suite of parsimonious mechanistic models that incorporate the effects of both school vacations and/or humidity and asses their ability to explain and forecast influenza incidence for small geographically contained populations (counties). We use the model set to reveal the contributions that each factor makes for each county over multiple years. We also combine these model results with several other model variants to produce a super-ensemble for each ILI profile. Finally, we use these results in a forecasting mode to demonstrate the value of such an approach at various phases during the influenza season.

## Methods

### Data

We obtained county-level data directly from each respective public health departments (Maricopa, AZ, San Diego, CA, Eastern, MO, Nashville-Davidson, TN, and Eastern, VA). For simplicity, and because we believe these counties are representative of their regional areas, we refer to these populations by their state names. These datasets were chosen based on: (1) their geographical diversity, allowing us to explore different climatological profiles; and (2) completeness, allowing us to examine multiple seasons. Estimates for the total population for each county were obtained from the US 2010 census data [19]. All ILI data collected were in the form of weekly reporting. To be consistent with week numbering and dates, we adopted the Morbidity and Mortality Weekly Report (MMWR) calendar. Thus, a week begins on Sunday and ends on Saturday, and week number 1 is the week containing the first Wednesday of the year. Any January days occurring before week one are considered part of the final week (52 or 53) of the previous year. See Supporting Information, S1 for details of auxiliary data used.

### Models

We used a deterministic SIR model (see Supporting Information, Text S2) with a constant background rate of clinical report (not driven by influenza infection). We determined the joint posterior distribution for the model parameters using a Metropolis-Hastings Markov chain Monte Carlo (MCMC) procedure [20]. For each county we simulated four MCMC chains each with 10^7^ steps and a burn time of 2.5x10^6^ steps (see below for effective sample sizes). At each step a new set of parameter values was sampled from a log-uniform distribution (the minimum and maximum allowed values for the parameters are summarized in Table S2). Using this set of candidate parameters, we generated a model incidence for the county and calculated the log-likelihood of the data. The values of the new and previous log-likelihood were used in the standard rejection method to determine if the move should be accepted or rejected. Our MCMC procedure had an adaptive step size which ensured an acceptance rate of 20-30%. The chains typically had an effective sample size in the 200-2000 range (depending on county profile and the parameter). The numerical fitting procedure is described in more detail in the Supplementary Materials (Text S2 and S3).

When modeling a specific ILI profile we considered each dataset to be independent and used a deterministic S-I-R compartmental model framework with a time dependent reproduction number *R*_0_(*t*):

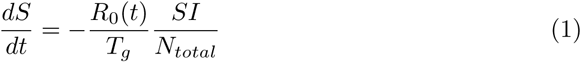

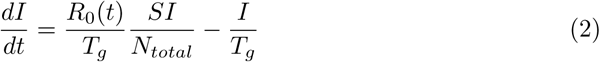

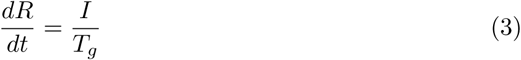

where *S* represents the number of susceptible individuals, *I* is the number of infectious individuals, *R* is the number of recovered individuals, and *N*_*total*_ = *S* + *I* + *R* is the total population. The time-parameter *t*_0_ is used to set initial conditions for the S-I-R equations were as follows:

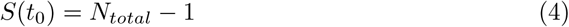

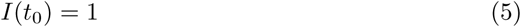

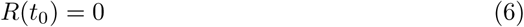

We extracted weekly incidence by integrating the rate that infections occurred,

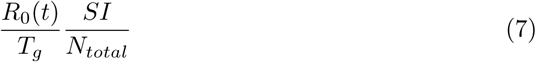

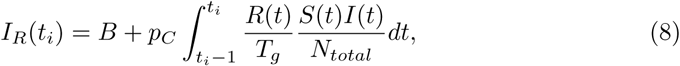

where *p*_*C*_ is the proportion of infectious people that present themselves to a clinic with ILI symptoms and *B* is a constant number of non-SIR cases or false-ILI. The integral runs over one week determining the number of model cases for week *t*_*i*_. This is how we relate the internal, continuous SIR model to the discrete weekly ILI incidence data. We describe the procedure used for fitting this property to the specific ILI profile in the Supporting Information, S2 Text.

In this study, we use four different time dependent models for the reproduction number, *R*_0_(*t*), as well as one time independent model. To achieve this, we write the transmission term in the most general way as a product of a constant reproduction number *R*_0*,B*_ and three time dependent terms:

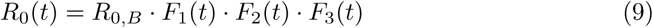

The first term *F*_1_(*t*), captures the dependence of the transmission rate on the specific humidity, the second on school vacation, and the third a simple two-value model [21].

Guided by [14], we define the effect of specific humidity as:

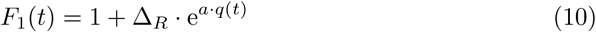

In contrast to [14], however, the values of the parameters *a* and Δ_*R*_ are fitted by the model. Δ_*R*_ must remain positive, and any effects of the specific humidity term can be deactivated by setting: Δ_*R*_ = 0. Specific humidity, *q*(*t*), is estimated using Phase-2 of the North American Land Data Assimilation System (NLDAS-2) database provided by NASA [22, 23]. The NLDAS-2 database provides hourly specific humidity (measured meters above the ground) for the continental US at a spatial grid of 0.125°, which we average to daily and weekly values.

For school vacations, we define:

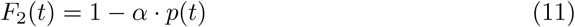

where *p*(*t*) is: 0 if the school is open; and 1 if the school is closed.

Data on school schedules were obtained directly from school districts within these counties. The effects of school vacations were explored by optimizing the parameter *α* over the range [0,1]. Higher magnitudes of *α* suggest that school closures cause a greater decrease to *R*_0_(*t*) and conversely small values of *α* indicate that the school schedule is not an important factor in transmission dynamics. As was the case for the specific humidity term, the effects of school vacations can be deactivated by setting *α* = 0. Alternately, the joint effects of humidity and school schedule can be explored by simultaneously optimizing the parameters of *F*_1_ (*t*) and *F*_2_ (*t*).

We also considered a model in which the underlying transmissibility could vary according to a step function, but for which the timing of the step and its amplitude were not informed by extrinsic factors such as school vacations and specific humidity. Rather, the step could be optimized to give the best fit to the data. For this “two-value” *R*_0_(*t*) model:

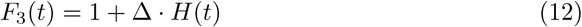

Where *H*(*t*) = 1 when *t*_*s*_ ≤ *t < t*_*f*_, and 0 otherwise.

We have found this model to be useful when modeling both military and civilian datasets [20, 21]. By allowing the parameter Δ to vary between −1 and +1, we can model both an increase and decrease in transmission due to behavior modification.

When *F*_1_(*t*), *F*_2_(*t*), and *F*_3_(*t*) are deactivated (Δ_*SH*_ = *α* = Δ = 0), the function reduces to a simple constant *R*_0_ model:

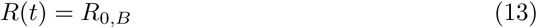

and the model is optimized with respect to only one parameter, *R*_0*,B*_.

In summary, we use five different models to describe the force of infection:

1. *R*_0_(*t*) depends on the specific humidity (model H)
2. *R*_0_(*t*) depends on the school vacation schedule (model V)
3. *R*_0_(*t*) depends on both specific humidity and the school vacation schedule (model HV)
4. *R*_0_(*t*) is constant (model NULL)
5. *R*_0_(*t*) is controlled by the two-value (step function) term *F*_3_(*t*) (model S)

Finally, the ensemble results (model E) are calculated as the unweighted average of these five model results.

The numerical aspects of the algorithm are discussed in more detail in Supporting Information S3.

## Model Performance Quantification

To compare the performance of various models, we introduce the Percentage of Deviance Explained (PDE)

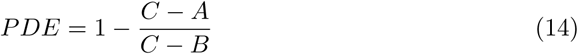

where *A* is the log-likelihood of the model we are evaluating, *B* is the log-likelihood of the NULL model, and *C* is the saturated log-likelihood of the data. Thus PDE describes the percentage of the deviance explained by model *A* relative to that explained by a naive SIR model (NULL). When calculating likelihoods, we assumed that the value obtained from the model was the mean of a Poisson distribution from which the data arose.

In the context of forecast performance evaluation, there is no guarantee that the individual models will perform as well or better than the NULL model. Thus it is possible to generate negative PDE values, which simply means that the forecast for the model in question performed worse than the NULL model forecast.

## Results

There was substantial variability in the data for both reported cases and potential drivers of transmissibility (Fig 1, top). Across all five populations, peak weekly-cases-reported varied by at least 400% over the study period. The duration of individual epidemics also varied, with the width of the curve at half maximum ranging between three and 11 weeks. Both specific humidity and school vacation schedule varied across the period of the study in the five different populations (Fig 1, bottom). The annual trend in specific humidity was reasonably consistent, but with significant, short-timescale fluctuations. School vacation times were similar amongst the different populations, but with some systematic differences. The start of the winter holiday vacation, in particular, often differed by a full week.

**Figure 1.**
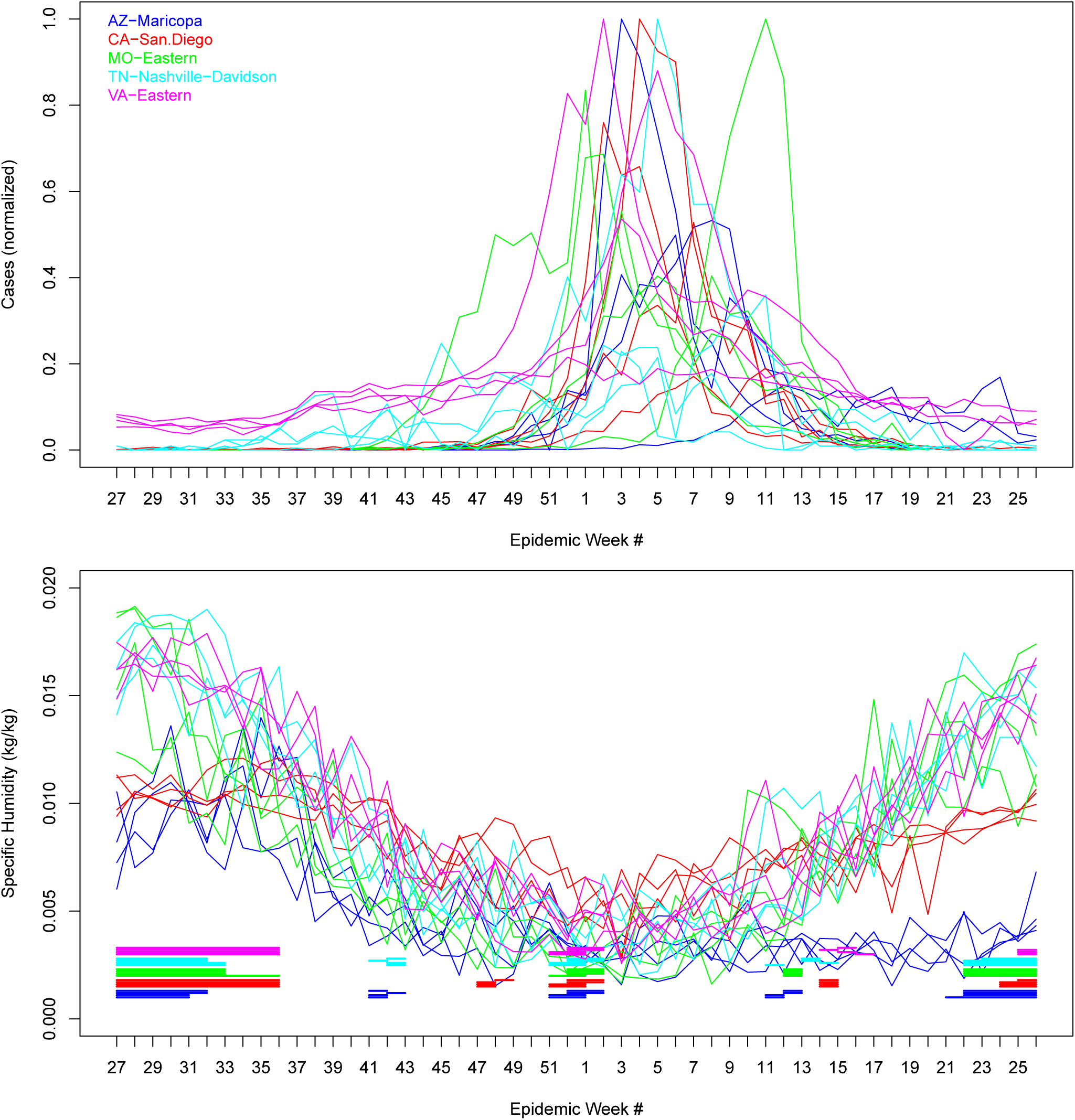
(Top) Incidence profiles for five counties during four seasons as a function of epidemic week number. The counties are delineated by different colors. (Bottom) Specific humidity as a function of epidemic week number for each season and county shown in the top panel. The times of school vacations are shown by the horizontal bars at the base of the plot. Each season’s vacation interval for a particular county is stacked upon the earlier one.

Using a model that contains terms for both specific humidity and school vacations (Model HV, see methods) we fitted incidence time series for all locations and all years (Fig 2). Assessed visually, the model was able to reproduce gross features of the epidemics such as peak height, width at half maximum and time of take-off. Interestingly, it was also able to reproduce some higher resolution features of incidence. For example, during the 2010-11 season, the incidence profile in AZ, CA, TN and VA flattened during the takeoff phase around the time of the school vacations.

**Figure 2.**
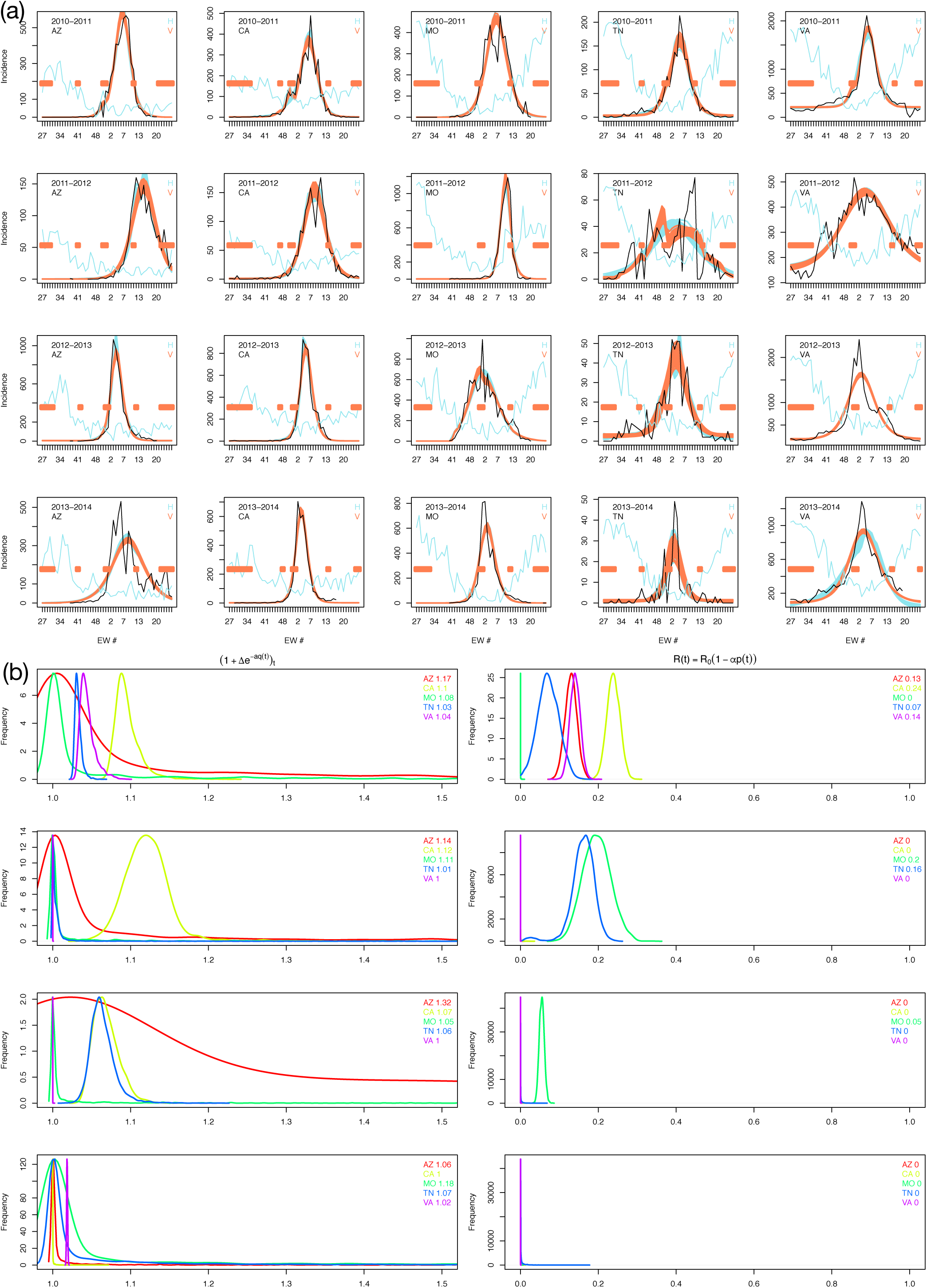
(A) Incidence profiles, and fits (using the specific humidity (H) and school vacation (V) models) for all five counties during four influenza seasons. Each of the five columns shows one county and each row corresponds to one season (see legend on top-left of each panel). In each panel the ILI incidence profile is shown in black and the results of the fits using the H and V models are shown in orange and blue, respectively. The jagged blue line denotes the weekly averaged specific humidity and the horizontal orange bars denote school vacations. The fits are 1,000 randomly chosen trajectories from the second half of the MCMC chain (which has 1 ×10^7^ steps.) (B) Left column: The poster<u>ior</u> distribution of the specific humidity term, using the yearly average of humidity: 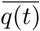 and the two parameters Δ_*R*_ and *a* that control the effect of humidity on *R*_0_(*t*). Top to bottom: 2010-2011, 2011-2012, 2012-2013, and 2013-2014 seasons. For each season we denote the five different counties with five different colors (see legend on top-right of each panel). The mean value of this term is shown in the top right of each panel. For Maricopa county in Arizona (red curves) this term deviates from zero for all four seasons, for San Diego County in CA (yellow curves) for three out of the four, for Eastern Missouri (green curves) for all four seasons, for Nashville-Davidson county in Tennessee (blue curves) for three out of four seasons and for Eastern Virginia (purple) for two (or possibly three) of the four seasons. Right column: The posterior distribution of the parameter *α* that controls the effect of school vacation on *R*_0_(*t*). See Eq (11) in text for more details. The mean value of *α* is shown in the top-right of each panel.

In our parameter estimates, these high resolution patterns were reflected in the posterior densities of the parameters that govern school vacations and specific humidity. We found evidence that both specific humidity and school vacations were important during some years in explaining the data but not during others, and sometimes both made contributions. For example, in Arizona and California, humidity appears to play a role in three out of the four seasons. In Tennessee, humidity is consequential in two seasons, while for Missouri and Virginia, the term was only significant in one season. The school vacation schedule appeared to be important only in one or two of the seasons across these populations.

Although the impact of each factor is not uniform across years and populations, parameter estimates that govern these features either take their null value or are bound within a narrow range (Fig 2(b)). Point estimates for the school vacations parameter cluster at either 0 or around 0.2. Similarly, the factor that governs the contribution of specific humidity seems limited to be below 1.15. Thus, when humidity is important, it appears to increase *R*_0_(*t*) by approximately 10-15%, while school vacations, when important, can cause *R*_0_(*t*) to decrease by up to 20%.

Further support for a causal relationship between specific humidity and transmission, albeit tentative, comes from a comparison of the humidity traces in Fig 2(a) and the amplitudes of the specific humidity transmission term in Fig 2(b) (left column). The locations MO, TN, and VA show deeper “U” shaped specific humidity profiles compared with AZ and CA, with the former group also showing a significant humidity effect. However, no apparent “threshold” for humidity appears to be present.

We used the constant *R*_0_(*t*) model (Model 4) as a NULL model to compare the performance of the other models across years and populations, again by fitting to the entire epidemic (Methods, Fig 3). Model S, the model with a step change in underlying transmissibility (Methods, Eq (12)) was always able to explain more of the deviance between the data and the NULL model than were other model variants (Table S1). However, the ranking and explanatory power of the different models (driven by school vacations and specific humidity) varied by population and by year. For some population-year combinations, either school vacations alone (Model V) or specific humidity alone (Model H) achieved similar explanatory power to the more flexible variable transmissibility (Model S).

**Figure 3.**
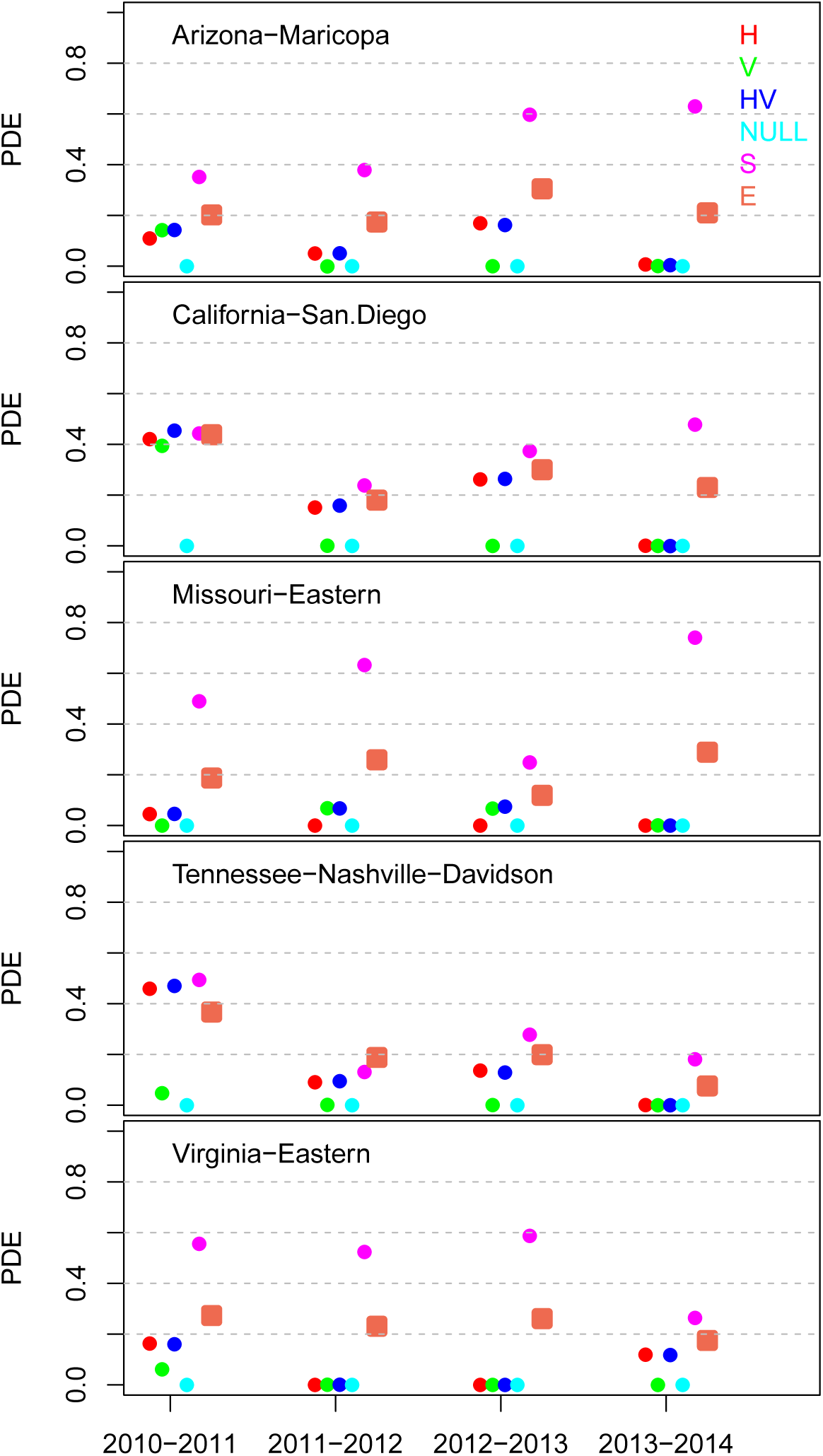
Percentage of deviance explained (PDE) for each season/model and for all five counties. The PDE is defined as: 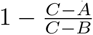, where *B* and *C* are the likelihoods of the NULL model and data respectively. *A* is the likelihood of each of the five models as well as the ensemble model (which is calculated as the average of the five models). By definition the PDE is zero for the NULL model which we have chosen to be the fixed *R*_0_(*t*) model and cannot exceed one.

We assessed the forecasting accuracy of individual models (H, V, HV, S) for all future time points, again, relative to the simple NULL SIR model. Our single measure of overall accuracy was the proportion of future deviance explained at any point in time, with deviance defined relative to the SIR model fit at that time point (Figs 4, 5, S4). Our results suggest that models including school vacation and specific humidity have the potential to increase forecast accuracy, but that gains in accuracy for any given model are likely to be location-specific and may also depend on the phase of the epidemic relative to its peak. School vacation terms and specific humidity terms consistently improved forecast accuracy for locations in California and Tennessee, when incorporated as Models V, H or HV. The step model, Model S, appeared to improve overall forecast accuracy in the second half of the season across all locations. A naive ensemble model in which all the individual models were given equal weight did not obviously improve performance over Models HV or S in those regions and during those periods where models HV and S appeared to be effective.

**Figure 4.**
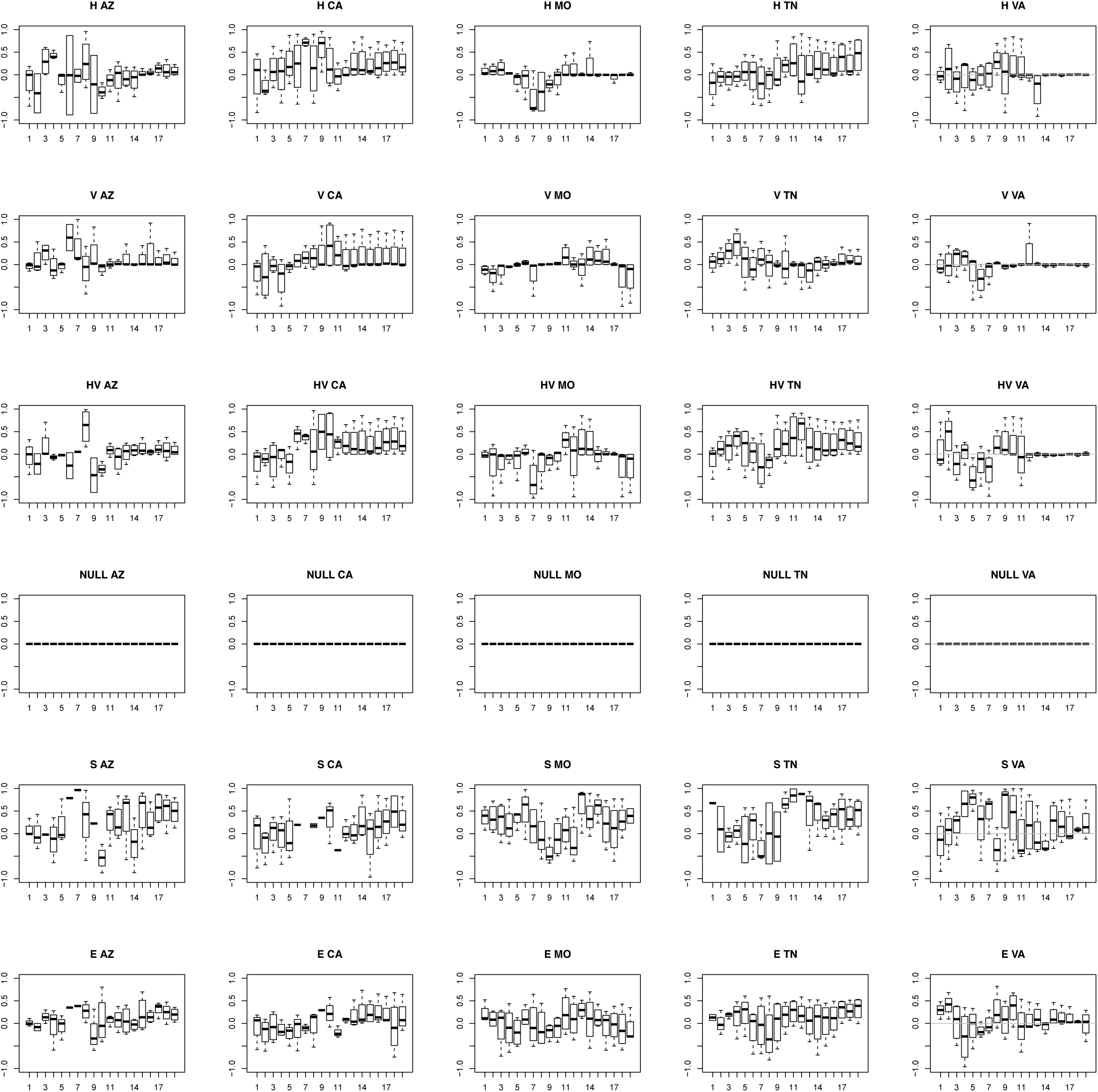
Forecast PDE as a function of epidemic week number. Here, we have fit each model to incidence data up to and including the epidemic week. The PDE then relates the resulting forecast to incidence for the remainder of the season. Each column is one of the five counties and each row corresponds to a model (H-specific humidity, V-school vacations, HV-specific humidity and school vacations, NULL-constant *R*_0_(*t*), S-two-value step function, E-ensemble). The PDE for each county/model is defined as: 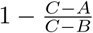. *A*, *B* and *C* are the likelihoods of the model, the NULL model and the data, respectively. The NULL model is that of fixed *R*_0_(*t*), and its constant PDE value of zero is marked in all the panels by the horizontal grey dashed line.

**Figure 5.**
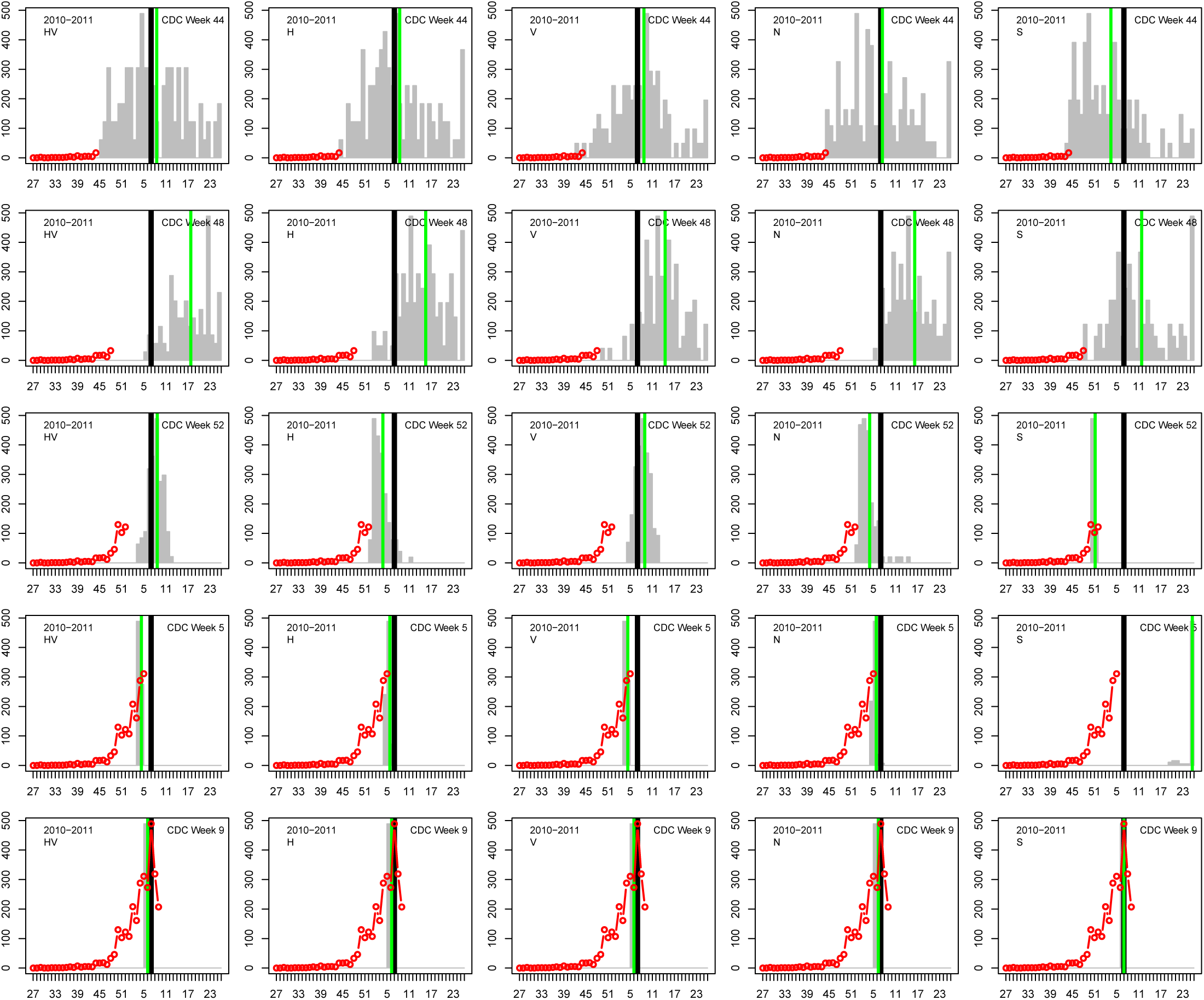
Comparison of each of the five models’ ability to predict the timing of the peak of the influenza season for San Diego County for the 2010-2011 season. In each column (from left to right) we show the results of one of the models: HV, H, V, fixed *R*_0_(*t*) and the two-value *R*_0_(*t*). The gray bars are the distribution of predicted peak weeks resulting from the MCMC chain (using 1,000 randomly selected trajectories), the green bar is the mean value and the black bar is the observed peak week. The red line/circles denote the data used to make the prediction. From top to bottom: epidemic week 44, 48, 52, 5 and 9.

As an example of a specific feature of epidemic incidence, we also compared the ability of the different models to predict the peak week by fitting them to data from only the early weeks of each influenza season (Figs 5 and S3). Early in the season, none of the models were able to predict the timing of the peak with any accuracy. However, after case data showed a clear rising pattern that could be fit to exponential growth, models V, H and HV (i.e.,s school vacations, specific humidity and both school vacations and specific humidity) were able to predict the timing of the peak; albeit with clear season-dependent structural biases. The additional parametric freedom of Model S actually reduced its ability to predict the timing of the peak.

## Discussion

In this study, we have generated evidence that both school vacations and specific humidity can have an impact on the incidence of influenza for small populations within the US when appropriately flexible models are fit that include information about these two mechanisms. Further, we have used a retrospective forecasting approach to show that those same models may have better forecast accuracy than a simple SIR-like model that does not include information on school vacations and specific humidity. We also developed a “super-ensemble;” However, we did not find evidence that this model performed any better – in either explanatory mode or forecast mode – than a single model that permitted the use of auxiliary data from both school vacations and specific humidity.

Several studies have estimated the reduction in transmissibility during school closures, both during seasonal periods [13] as well as during pandemics [13]. More recently, [24] showed that school vacations delay epidemic peaks and act to synchronize incidence profiles at different locations. The effects of humidity in modulating influenza transmission have also been well studied [8, 14, 25], including the benefits of ensemble models incorporating specific humidity, which could be used to provide forecasts in real-time [9]. By obtaining both humidity data and school vacation data for the same small populations, and examining their effect within a mechanistic model, we extend these previous studies by describing a complex system where different components are important at different times.

Our results are broadly consistent with the work of [26] in which both school closures and specific humidity were both considered. They estimated the impact of both of these processes during the 2009 influenza pandemic in Mexican states, finding that the the spatial structure of the pandemic could be explained by a combination of factors: high specific humidity on some states driving activity, and school vacations during the summer preventing further transmission. Additionally, they attributed anomalously large outbreaks in some states to differences in residual susceptibility (a factor not likely significant for seasonal influenza). However, we note that [26] was implemented at a much larger spatial scale than we have used here and that the fine detail of state-level incidence did not appear to be driven by either school vacations or specific humidity, nor was the forecast potential of the model explored.

While it is encouraging that the model results indicate a role for school vacations and humidity, it is not clear why the results were not more consistent. There is a risk that we have over-fit the models and thus overstated the importance of school vacations and specific humidity. We suggest that the relationship between epidemic onset time and the start of the vacations is crucial for assessing whether vacations are going to play a role, and this type of prior information can be convolved into a model prediction. Perhaps more data from multiple populations will reveal that only when the vacation occurs during the early exponential rise do they cause the profile to stall for several weeks, and, in turn, delay the arrival of the peak. If it occurs too late (at, or after the peak), the contribution of schools may be substantially weakened. Similarly, we suggest that humidity only plays a role in locations where the humidity shows significant variation. Moreover, the phasing of the variation must be such that it can act as a catalyst for the outbreak, in essence, creating the “spark” that drives a steeper rise in the ILI incidence profile.

We did not include age-classes explicitly in this study, largely because this information is not available in the datasets constructed from the county office weekly reports. Were age-stratified data available, we would certainly have refined our models to take this into account. However, although it is likely that the detailed epidemic dynamics we observe in our study populations were influenced by age effects, particularly between school-age children and adults, the non-age structured model we used likely performs well as an average description of the epidemic.

These results suggest that many different models will need to be triaged and tested for each time point in each population and then only models for which there is credible support be included in final forecasts. We note that even though we did not find our simple ensemble approach to be an improvement over our flexible single model [20, 21], the adoption and development of more sophisticated ensemble methods [8, 9] may achieve exactly that goal. Given the relatively high variations in both school vacations and specific humidity across small spatial scales, future ensemble approaches may need to allow different model weights for different small units of geographical space.

### Disclaimer

The findings and conclusions in this report are those of the authors and do not necessarily represent the views of the Department of Health and Human Services or its components, the US Department of Defense, local country Ministries of Health, Agriculture, or Defense, or other contributing network partner organizations. Mention of any commercial product does not imply DoD endorsement or recommendation for or against the use of any such product. No infringement on the rights of the holders of the registered trademarks is intended. No funding bodies had any role in study design, data collection and analysis, decision to publish, or preparation of the manuscript.

---

**Figure S1.**
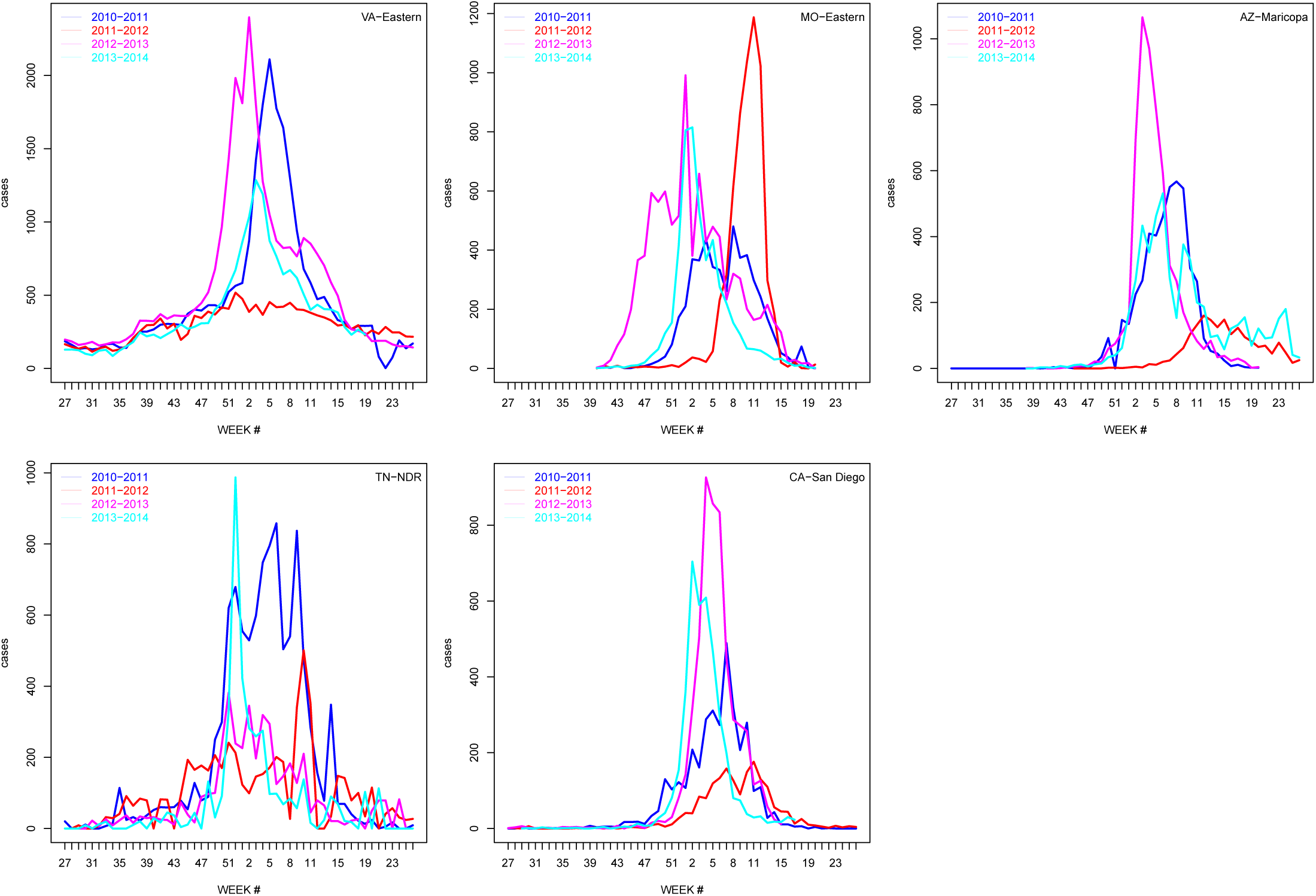
Incidence profiles for each season are shown for each of the five counties studied for the four influenza seasons.

**Figure S2.**
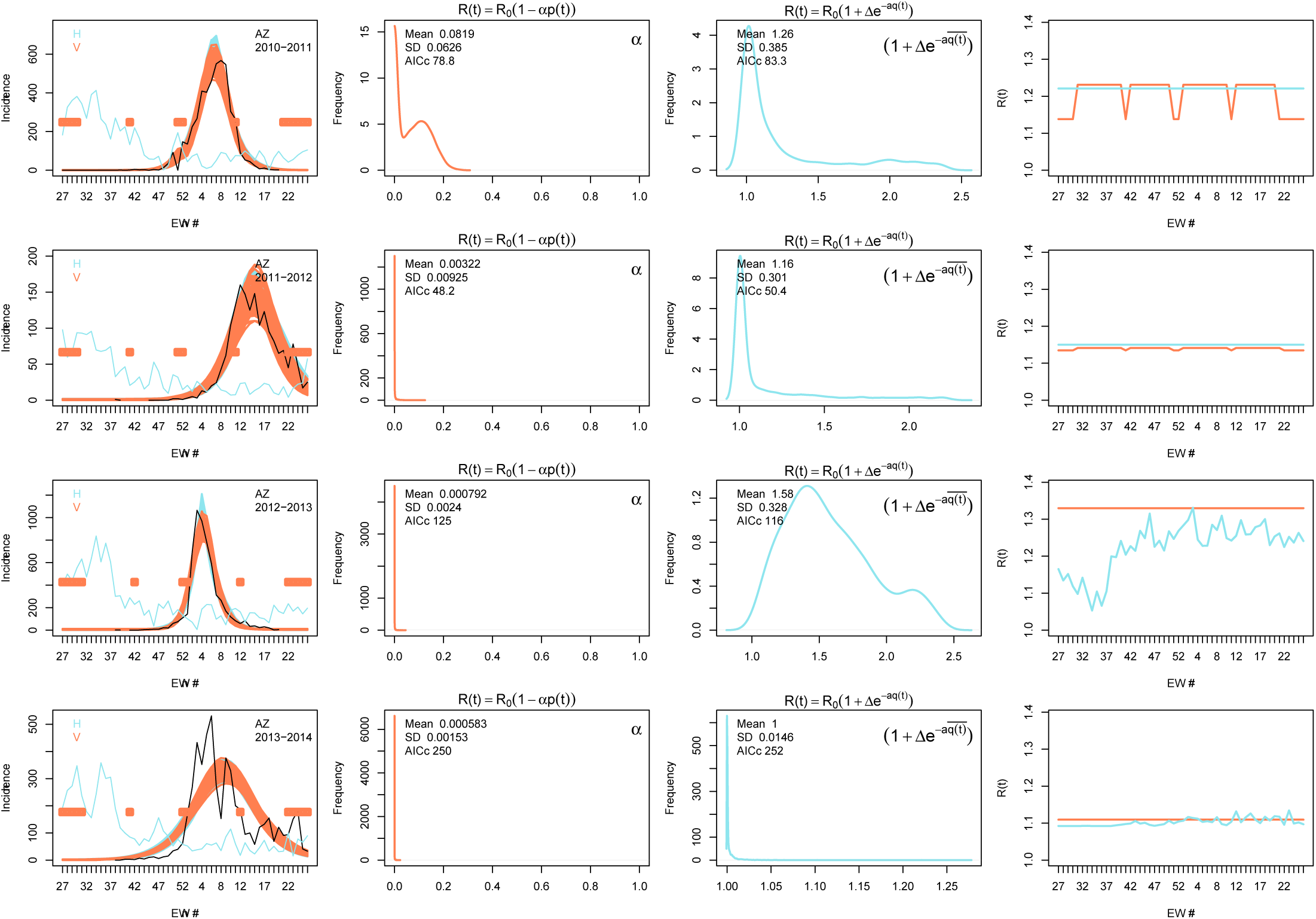
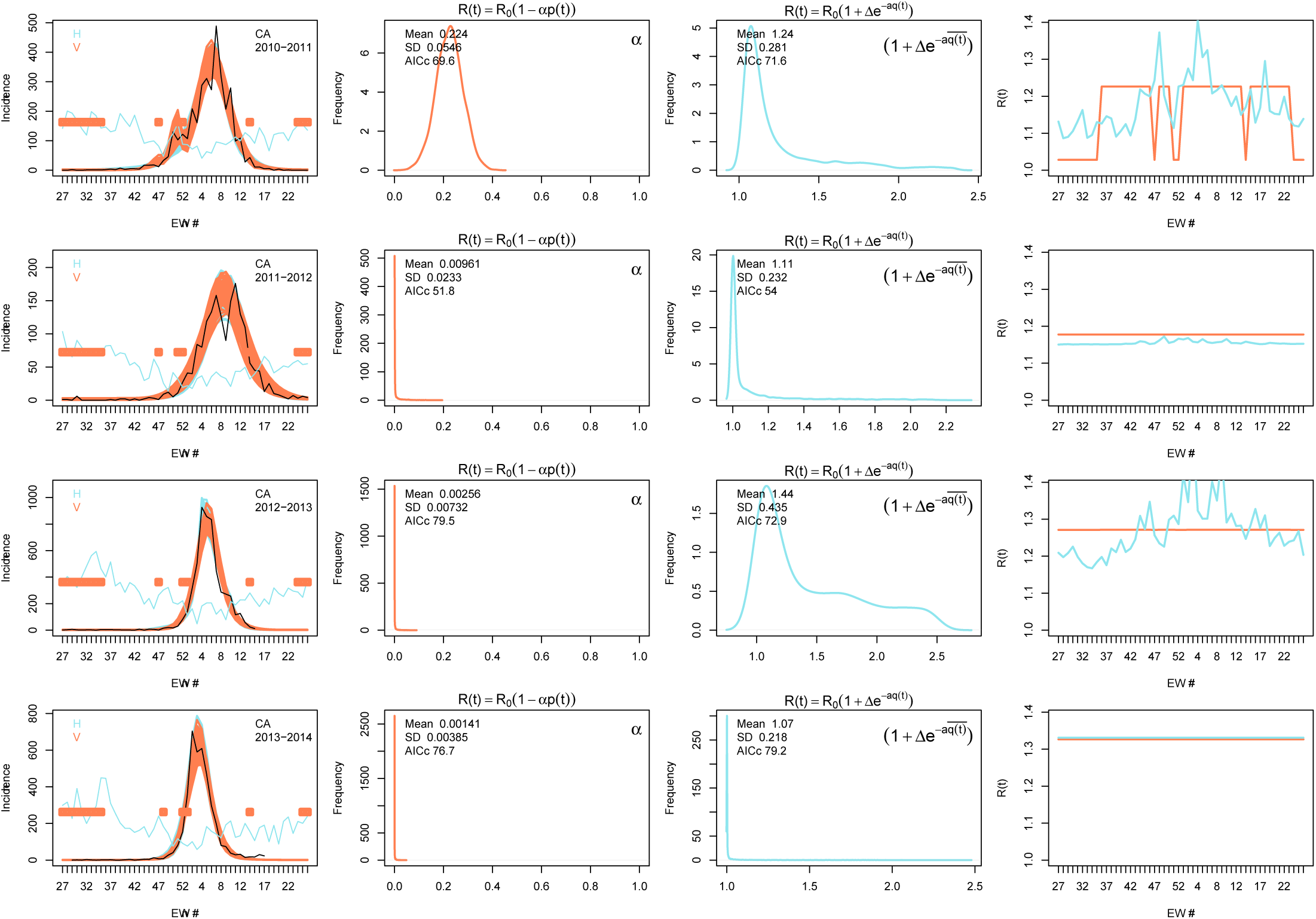
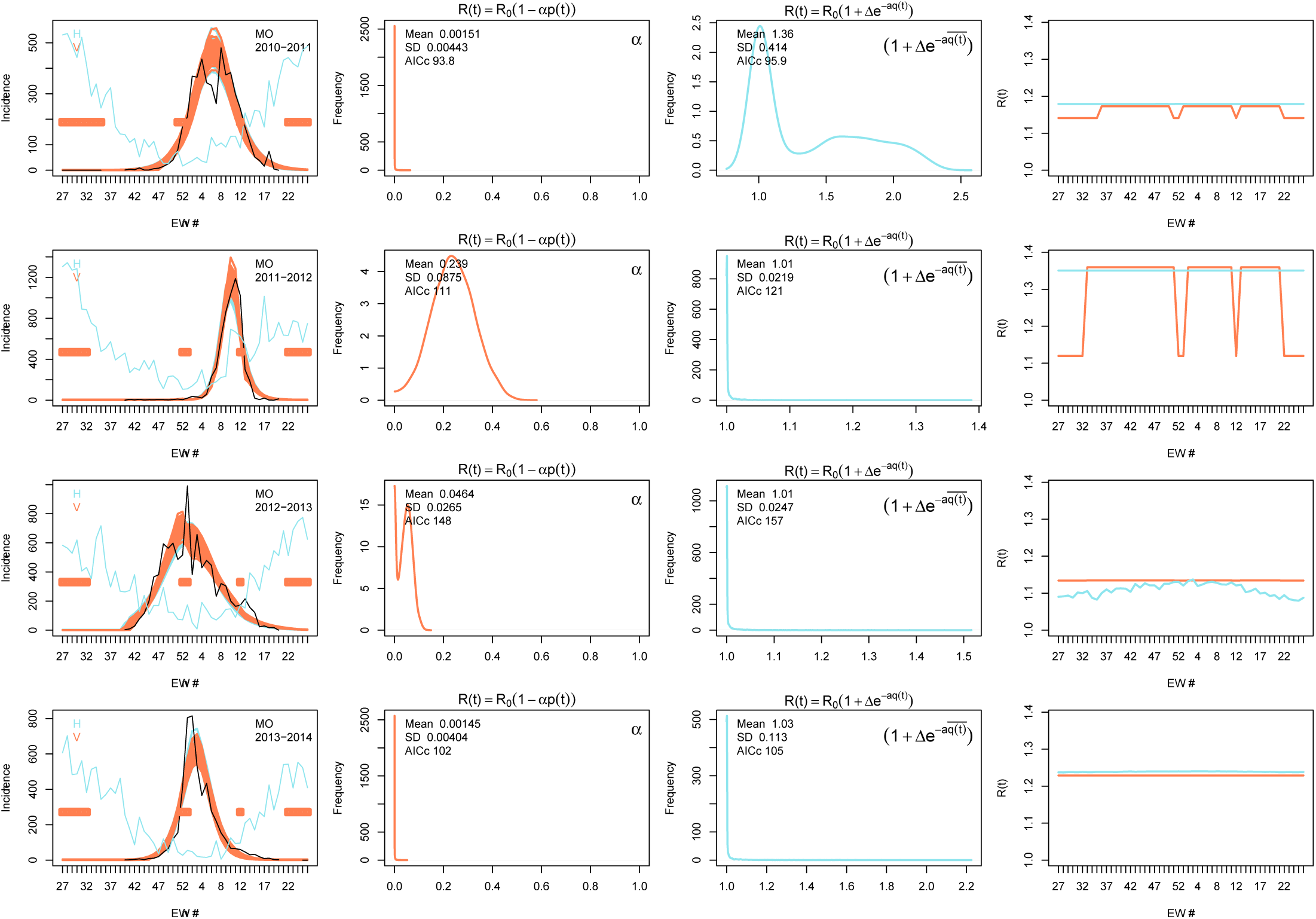
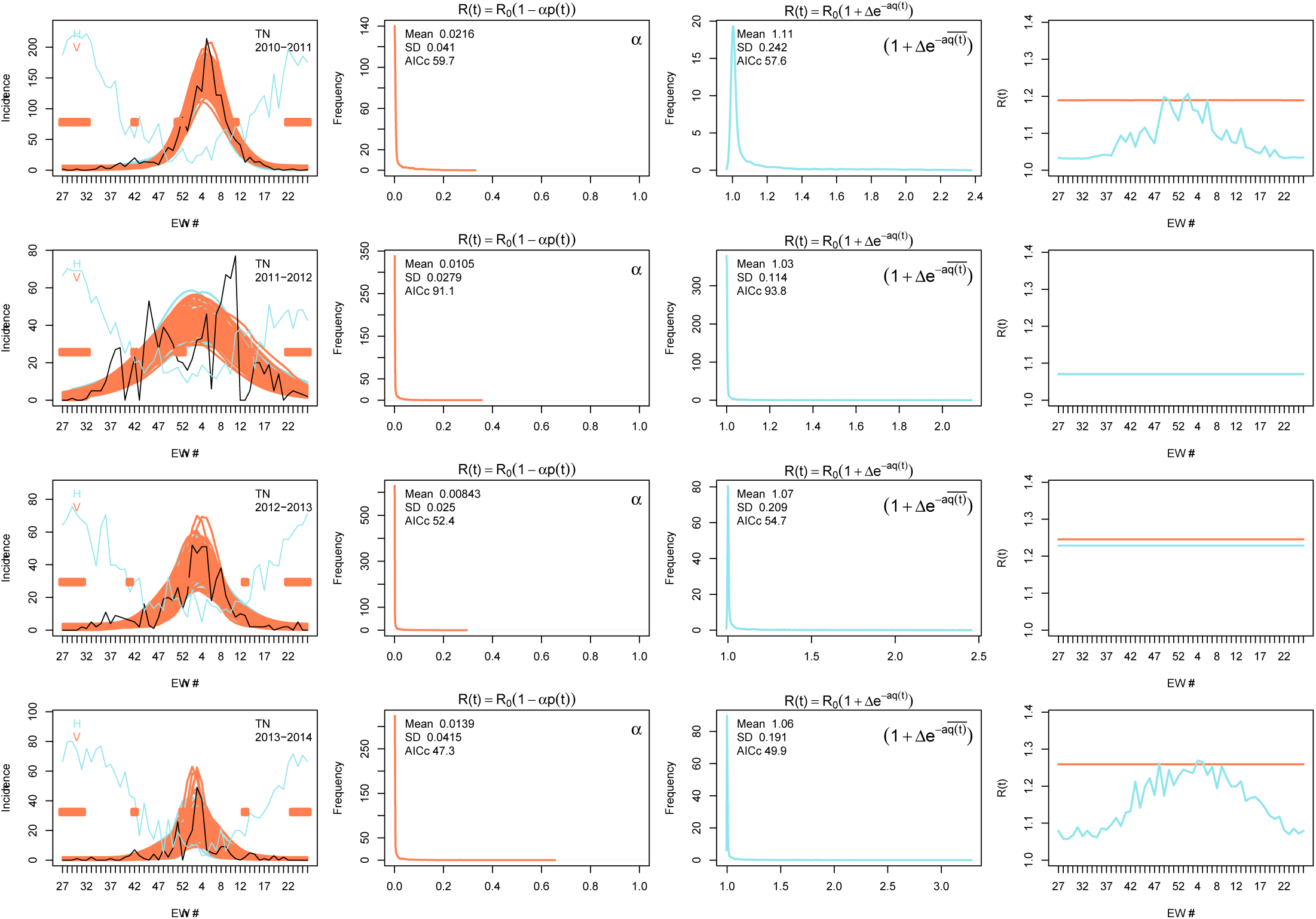
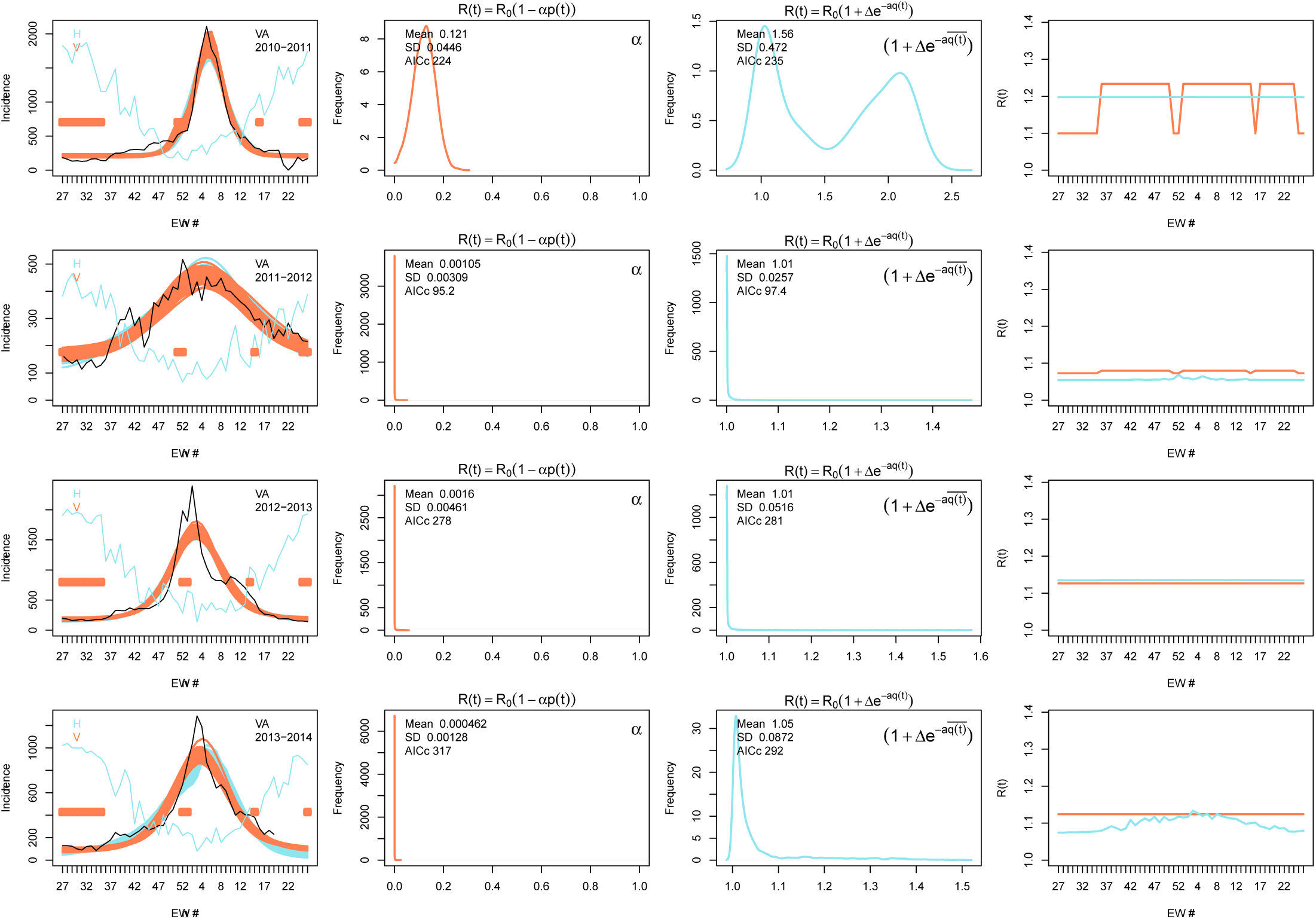
Influence of school vacations and specific humidity for each of the five regions studied for the four influenza seasons. Each of the five panels (A-E) is for one county and each row within a county is for a given influenza season (see legend on top right of each panel in the left column.) Left column: in each panel we show the ILI incidence profile in black and the results of the MCMC procedure (randomly selected 1,000 profiles from the second half of the MCMC chain) for the specific humidity and and school vacation models in blue and orange, respectively. The blue line denotes the specific humidity and the horizontal orange bars specify the timing of school vacations. Second column: The posterior distribution of the parameter *α* that controls the effect of the school vacation term. Third column: the posterior distribution of the specific humidity term calculated using the yearly average value of 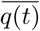. Fourth column: the time dependent force of infection, *R*_0_(*t*) as a function of epidemic week for the school vacation (orange) and specific humidity (blue) models. We randomly sample 1,000 trajectories from the second half of the MCMC chain and used the mean parameter values to generate the force of infection impact as a function of epidemic week.

**Figure S3.**
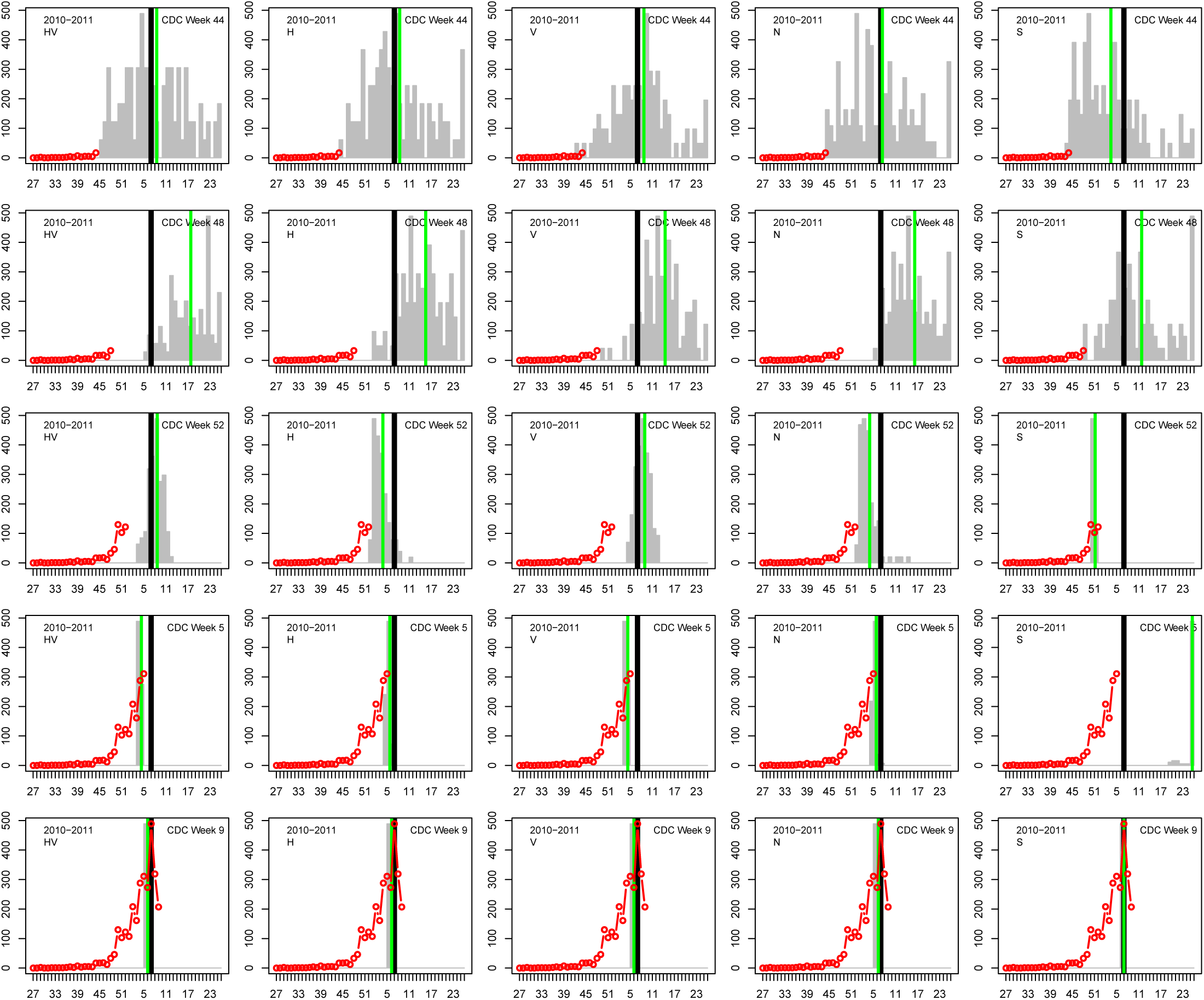
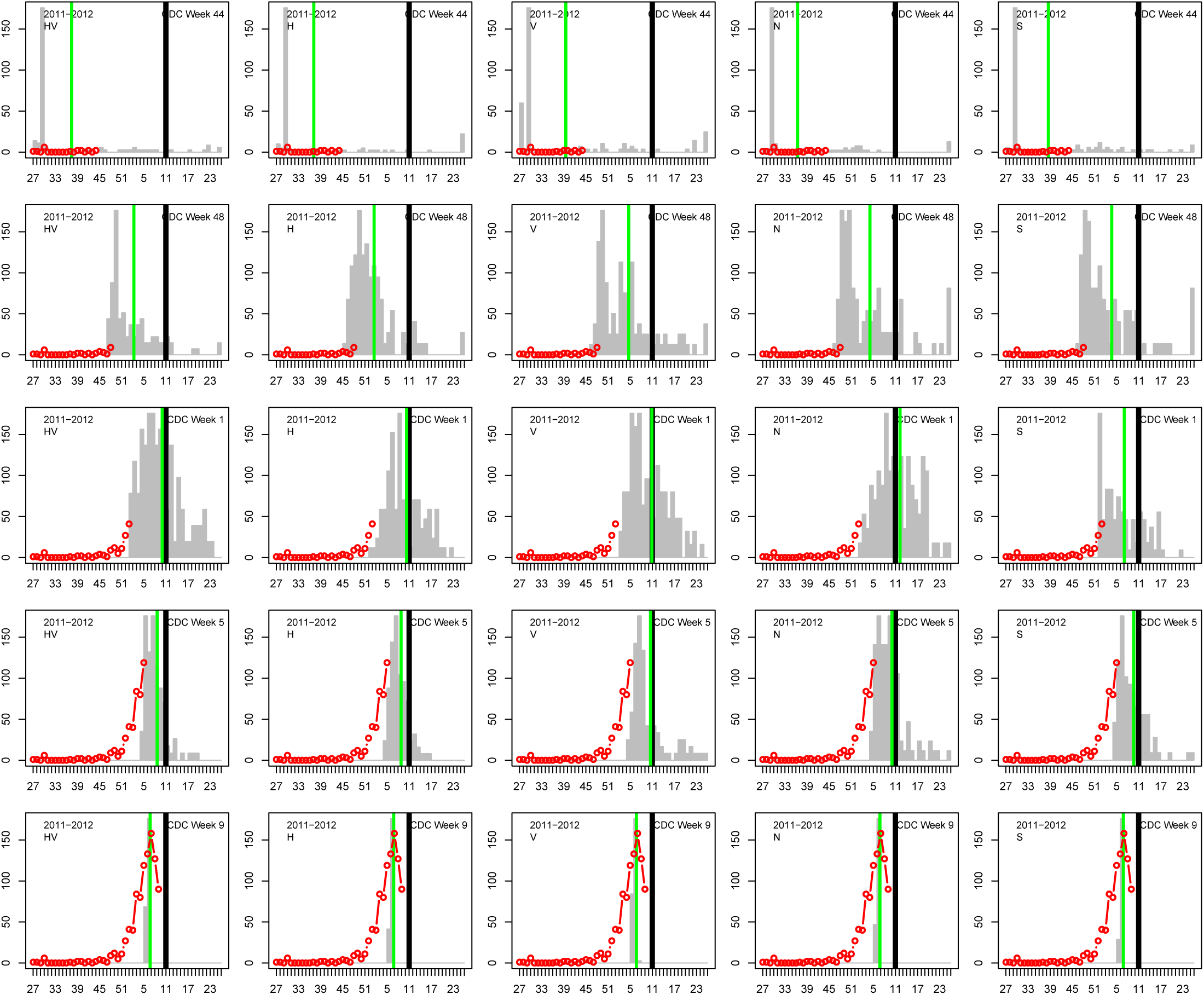
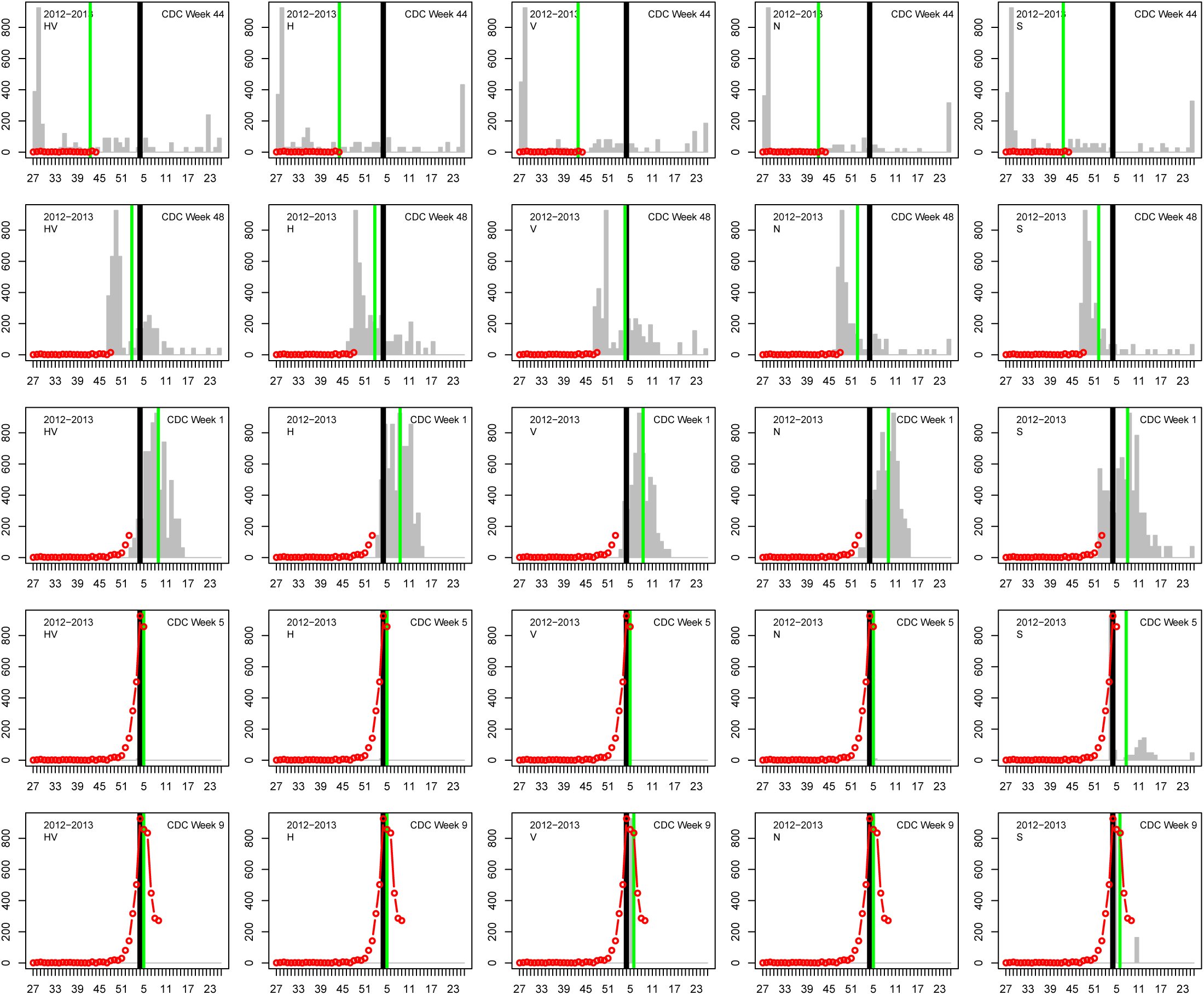
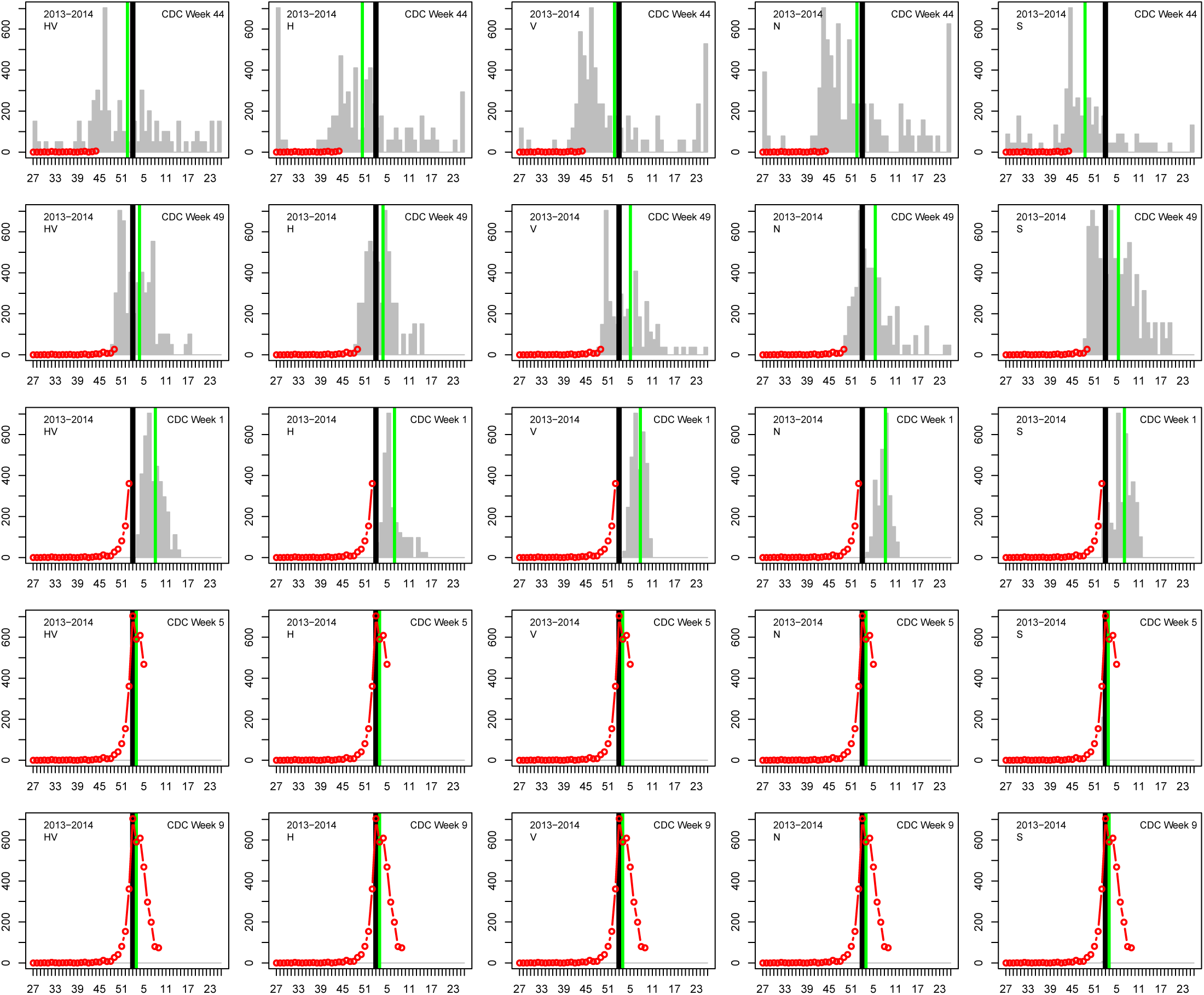
Comparison of each of the five models’ ability to predict the timing of the peak of the influenza season for San Diego County during each of the seasons from 2010 through 2014 (A-D). In each column (from left to right) we show the results of one of the models: HV, H, V, fixed *R*_0_(*t*) and the two-value *R*_0_(*t*). The grey bars are the distribution of predicted peak weeks resulting from the MCMC chain (using 1,000 randomly selected trajectories), the green bar is the mean value and the black bar is the observed peak week. The red line/circles denote the data used to make the prediction. From top to bottom: epidemic week 44, 48, 52, 5 and 9.

## Supporting information

### S1 Text. Auxiliary Data

The specific humidity *q*(*t*) for all continental U.S. locations is produced using the Phase-2 of the North American Land Data Assimilation System (NLDAS-2) [22, 23] database provided by NASA http://ldas.gsfc.nasa.gov/nldas/NLDAS2model.php/. The NLDAS-2 database provides hourly specific humidity (measured 2-meters above the ground) for the continental US at a spatial grid of 0.125° which we average to daily and weekly SH. The weekly data is then interpolated to the appropriate position. For populations outside the continental US (of which there were none in the current study), and for the states of Alaska and Hawaii, we obtain the specific humidity data from NOAA’s NCEP-NCAR Reanalysis project (see for example: http://iridl.ldeo.columbia.edu/SOURCES/.NOAA/.NCEP-NCAR/.CDAS-1/.DAILY/.Diagnostic/.above_ground/.qa/) which provides daily (again 2-meter above ground) specific humidity data on a spatial grid of 2.5° for the entire world. This data is averaged and interpolated using the same procedure as for the NLDAS-2 dataset. The specific humidity database is continually updated using these two dataset sources.

School vacation data were collected from a representative school district in each county. Each week of the school year was determined to be either ‘in session’ or ‘on vacation’ and assigned a value of 0 or 1 respectively. The week of Thanksgiving Day was always categorized as vacation. Otherwise, only weeks with three or more days of no classes were considered vacation weeks.

### S2 Text Numerical Techniques Employed in the Model

The model ILI fitting procedure determines the joint posterior distribution for the model parameters (determined by the user-selected *R*_0_(*t*) model) using a Metropolis-Hastings Markov Chain Monte Carlo (MCMC) procedure. After the user selects which parameters to optimize and which to not, and what values to set the latter to, the user must specify the number of MCMC chains and steps per chain. The model then randomly initializes the parameters that are optimized (using a log-uniform distribution for all the parameters except those that can be negative), integrates the coupled S-I-R and influenza incidence equations, and generates a candidate ILI profile. The goodness of fit is measured by the Akaike Information Criterion (AIC), which is a measure of the relative goodness of fit of a model:

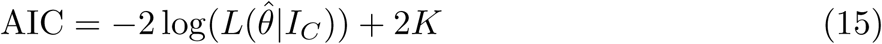

where 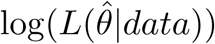 is the value of the maximized log-likelihood over the model parameters (*θ*), given the observed cases *I*_*C*_. When the total number of parameters (*K*) is large relative to the sample size (*n*), the reduced Akaike Information Criterion is preferred:

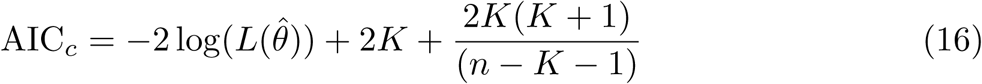

which is what the model uses in this study. The log-likelihood stems from a Poisson probability density

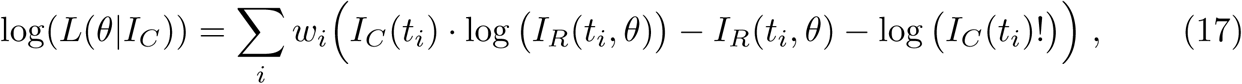

where *I*_*R*_(*t*_*i*_, *θ*) is the model point generated for week *t*_*i*_ as a result of parameter set *θ*. Additionally, the vector *w* has been added to allow the user to vary the weight given to each weekly data point. To maximize log(*L*(*θ* | *I*_*C*_)), the value of this likelihood is compared to a new likelihood calculated using a set of randomly displaced parameters in a standard rejection method to determine if the move is accepted or rejected. This MCMC procedure is executed as many times as the user has defined (this is the chain’s length mentioned above) and the model keeps the history of the chain parameters and *AIC*_*c*_ values. Once a chain is completed, it’s history statistics and results are summarized and written to tables (csv format), a binary RData file and pdf/png plots. The model MCMC chains have an adaptive step size which results in an acceptance rate of 20% – 30%.

**Table S1.** A summary of Models/Parameters used in this study. As described in the main text, there are a number of ways to define *R*_0_(*t*). Table S1 contains a brief description of each model and specifies which parameters are being optimized. Parameters that are not being optimized are generally set to their default value.

| Model # | Description | Optimized Parameters |
| --- | --- | --- |
| 1 | Specific Humidity only | $R_0, \Delta_R, a, t_0, B, pC$ |
| 2 | School only | $R_0, t_0, B, pC, \alpha$ |
| 3 | School and specific humidity terms | $R_0, \Delta_R, a, t_0, B, pC, \alpha$ |
| 4 | Constant $R_0$ | $R_0, t_0, B, pC$ |
| 5 | Two-step $R_0$ (see eq (12)) | $R_0, t_0, B, pC, \Delta, t_s, \Delta t$ |

**Table S2.** Summary of parameters. The “Optimum Range” is the range of values that the optimizer is allowed to vary over. For the parameters that control *R*_0_(*t*), their default value is often 0. Thus when a parameter is not being optimized, it is set to zero which deactivates that term. Both the default and range of values for the baseline parameter *B* depend on *B*_*est*_. For a single influenza year (e.g., Summer 2010–Summer 2011), *B*_*est*_ is the average of the first five and last five weeks of ILI data.

| Par | Description | Default | Optim Range |
| --- | --- | --- | --- |
| $R_0$ | Baseline reproduction number | 1.4 | 0.5–4.0 |
| $T_g$ | Average infectious period (weeks) | 2.6/7 | (1/7)–1 |
| $t_0$ | Infection start week ($I(t_0) = 1$) | 1.0 | 1.0–40.0 |
| $B$ | Baseline: mag. of non-SIR ILI data | $B_{est}$ | $(0.1 * B_{est}) - (10 * B_{est})$ |
| $pC$ | Percent clinical | 0.01 | $10^{-6}$–1.0 |
| $\alpha$ | Reduction in $R_0$ for school closed | 0 | $10^{-6}$–1.0 |
| $\Delta_R$ | Max change in $R_0$ from specific humidity | 0 | $10^{-3}$–2.0 |
| $a$ | Exponential for specific humidity $R_0$-term | 200 | 1.0–500.0 |
| $\Delta$ | Increase/Decrease in $R_0$ (model 5) | 0 | -1.0–1.0 |
| $t_s$ | Start time of change in $R_0$ (model 5) | 0 | 1.0–40.0 |
| $dur$ | time duration: $t_f = t_s + dur$ (model 5) | 0 | 1.0–40.0 |

## References

1. Anderson RM, May RM. Infectious disease of humans: dynamics and control. Oxford Science Publications. 1992;.

2. Chis Ster I, Ferguson NM. Transmission parameters of the 2001 foot and mouth epidemic in Great Britain. PLoS ONE. 2006;.

3. Ferguson NM, Donnelly CA, Anderson RM. Transmission intensity and impact of control policies on the foot and mouth epidemic in Great Britain. Nature. 2001;.

4. Ong JBS, Mark I, Chen C, Cook AR, Lee HC, Lee VJ, et al. Real-time epidemic monitoring and forecasting of H1N1-2009 using influenza-like illness from general practice and family doctor clinics in Singapore. PloS one. 2010;5(4):e10036.

5. Chretien JP, Riley S, George DB. Mathematical modeling of the West Africa Ebola epidemic. Elife. 2015;4:e09186.

6. Perkins A, Siraj A, Ruktanonchai CW, Kraemer M, Tatem A. Model-based projections of Zika virus infections in childbearing women in the Americas. BioRxiv. 2016; p. 039610.

7. Biggerstaff M, Alper D, Dredze M, Fox S, Fung ICH, Hickmann KS, et al. Results from the centers for disease control and prevention’s predict the 2013–2014 Influenza Season Challenge. BMC Infectious Diseases. 2016;16(1):357.

8. Shaman J, Karspeck A. Forecasting seasonal outbreaks of influenza. Proceedings of the National Academy of Sciences. 2012;109(50):20425–20430. doi:10.1073/pnas.1208772109.

9. Shaman J, Karspeck A, Yang W, Tamerius J, Lipsitch M. Real-time influenza forecasts during the 2012–2013 season. Nature communications.2013;4.

10. Zhang X, Meltzer MI, Wortley PM. FluSurge—a tool to estimate demand for hospital services during the next pandemic influenza. Medical Decision Making. 2006;26(6):617–623.

11. Matheny J, Toner E, Waldhorn R. Financial effects of an influenza pandemic on US hospitals. Journal of Health Care Finance. 2007;34(1):58.

12. Cowling BJ, Lau EHY, Lam CLH, Cheng CKY, Kovar J, Chan KH, et al. Effects of school closures, 2008 winter influenza season, Hong Kong. Emerging Infect Dis. 2008;.

13. Cauchemez S, Valleron AJ, Boëlle PY, Flahault A, Ferguson NM. Estimating the impact of school closure on influenza transmission from Sentinel data. Nature. 2008;452(7188):750.

14. Shaman J, Pitzer VE, Viboud C, Grenfell BT, Lipsitch M. Absolute humidity and the seasonal onset of influenza in the continental United States. PLoS biology. 2010;8(2):e1000316+. doi:10.1371/journal.pbio.1000316.

15. Lowen AC, Mubareka S, Steel J, Palese P. Influenza virus transmission is dependent on relative humidity and temperature. PLoS Pathog. 2007;3(10):e151.

16. Viboud C, Alonso WJ, Simonsen L. Influenza in tropical regions. PLoS Med. 2006;3(4):e89.

17. Tellier R. Review of Aerosol Transmission of Influenza A Virus. Emerging Infectious Disease. 2006;12(11).

18. Lowen AC, Steel J, Mubareka S, Palese P. High temperature (30 C) blocks aerosol but not contact transmission of influenza virus. Journal of virology. 2008;82(11):5650–5652.

19. for New Media C, Promotions(C2PO). US Census Bureau 2010 Census; 2009. Available from: https://www.census.gov/2010census/data/.

20. Riley P, Ben-Nun M, Armenta R, Linker JA, Eick AA, Sanchez JL, et al. Multiple Estimates of Transmissibility for the 2009 Influenza Pandemic Based on Influenza-like-Illness Data from Small US Military Populations. PLoS computational biology. 2013;9(5):e1003064.

21. Riley P, Ben-Nun M, Linker JA, Cost AA, Sanchez JL, George D, et al. Early characterization of the severity and transmissibility of pandemic influenza using clinical episode data from multiple populations. PLoS Comput Biol. 2015;11(9):e1004392.

22. Xia Y, Mitchell K, Ek M, Sheffield J, Cosgrove B, Wood E, et al. Continental-scale water and energy flux analysis and validation for the North American Land Data Assimilation System project phase 2 (NLDAS-2): 1. Intercomparison and application of model products. Journal of Geophysical Research: Atmospheres. 2012;117(D3).

23. Xia Y, Mitchell K, Ek M, Cosgrove B, Sheffield J, Luo L, et al. Continental-scale water and energy flux analysis and validation for North American Land Data Assimilation System project phase 2 (NLDAS-2): 2. Validation of model-simulated streamflow. Journal of Geophysical Research: Atmospheres. 2012;117(D3).

24. Ewing A, Lee EC, Viboud C, Bansal S. Contact, travel, and transmission: The impact of winter holidays on influenza dynamics in the United States. Journal of Infectious Diseases. 2016; p. jiw642.

25. Shaman J, Kohn M. Absolute humidity modulates influenza survival, transmission, and seasonality. Proceedings of the National Academy of Sciences. 2009;106(9):3243–3248.

26. Tamerius J, Viboud C, Shaman J, Chowell G. Impact of school cycles and environmental forcing on the timing of pandemic influenza activity in Mexican States, May-December 2009. PLoS computational biology. 2015;11(8):e1004337.

